# Vms1p is a release factor for the Ribosome-associated Quality control Complex

**DOI:** 10.1101/301341

**Authors:** Olga Zurita Rendón, Eric K. Fredrickson, Conor J. Howard, Jonathan Van Vranken, Sarah Fogarty, Neal D. Tolley, Raghav Kalia, Beatriz A. Osuna, Peter S. Shen, Christopher P. Hill, Adam Frost, Jared Rutter

## Abstract

Eukaryotic cells employ the Ribosome-associated Quality control Complex (RQC) to maintain homeostasis despite defects that cause ribosomes to stall. The RQC comprises the E3 ubiquitin ligase Ltn1p, the ATPase Cdc48p, and the novel proteins Rqc1p and Rqc2p^1–3^. Following recognition and subunit splitting of stalled ribosomes, the RQC detects and assembles on 60S subunits that hold incomplete polypeptides linked to a tRNA (60S:peptidyl–tRNA)^4–8^. Ltn1p cooperates with Rqc1p to facilitate ubiquitination of the incomplete nascent chain, marking it for degradation^7,9,10^. Rqc2p stabilizes Ltn1p on the 60S^3–5,8^ and recruits charged tRNAs to the 60S to catalyze elongation of the nascent protein with Carboxy-terminal Alanine and Threonine extensions, or CAT tails, via a mechanism that is distinct from canonical translation^4,10^. CAT-tailing mobilizes and exposes lysine residues in the nascent chain, especially those stalled within the exit tunnel, thereby supporting efficient ubiquitination^10,11^. If the ubiquitin-proteasome system is overwhelmed or unavailable, CAT-tailed nascent chains aggregate in the cytosol or within organelles like the mitochondria^12–14^. Here we identify Vms1p as the tRNA hydrolase that releases nascent polypeptides for extraction and degradation in the RQC pathway.

---

Like other RQC components, Vms1p is conserved throughout Eukarya and promotes protein quality control in diverse settings. In S. *cerevisiae*, Vms1p localizes to mitochondria in response to mitochondrial stress or damage^15,16^. Mutants lacking Vms1p are sensitive to rapamycin, which impairs ribosomal protein synthesis^17,18^, although the mechanism for this sensitivity is unknown. We found that the *vms1*Δ strain is also sensitive to the protein synthesis inhibitor cycloheximide (CHX), as are other RQC mutants in a sensitizing *SKI* mutant background^11^. Surprisingly, deletion of any one of the RQC system components *RQC1*, *RQC2* or *LTN1*, was sufficient to reverse the lethality of the *vms1*Δ mutant in CHX (**Fig. 1a**). By contrast, deletion of the no-go decay^19,20^ components *DOM34* or *SKI7* had no effect. These data suggest that CHX causes accumulation of an RQC product that is toxic unless Vms1p is available to detoxify it.

**Figure 1.**
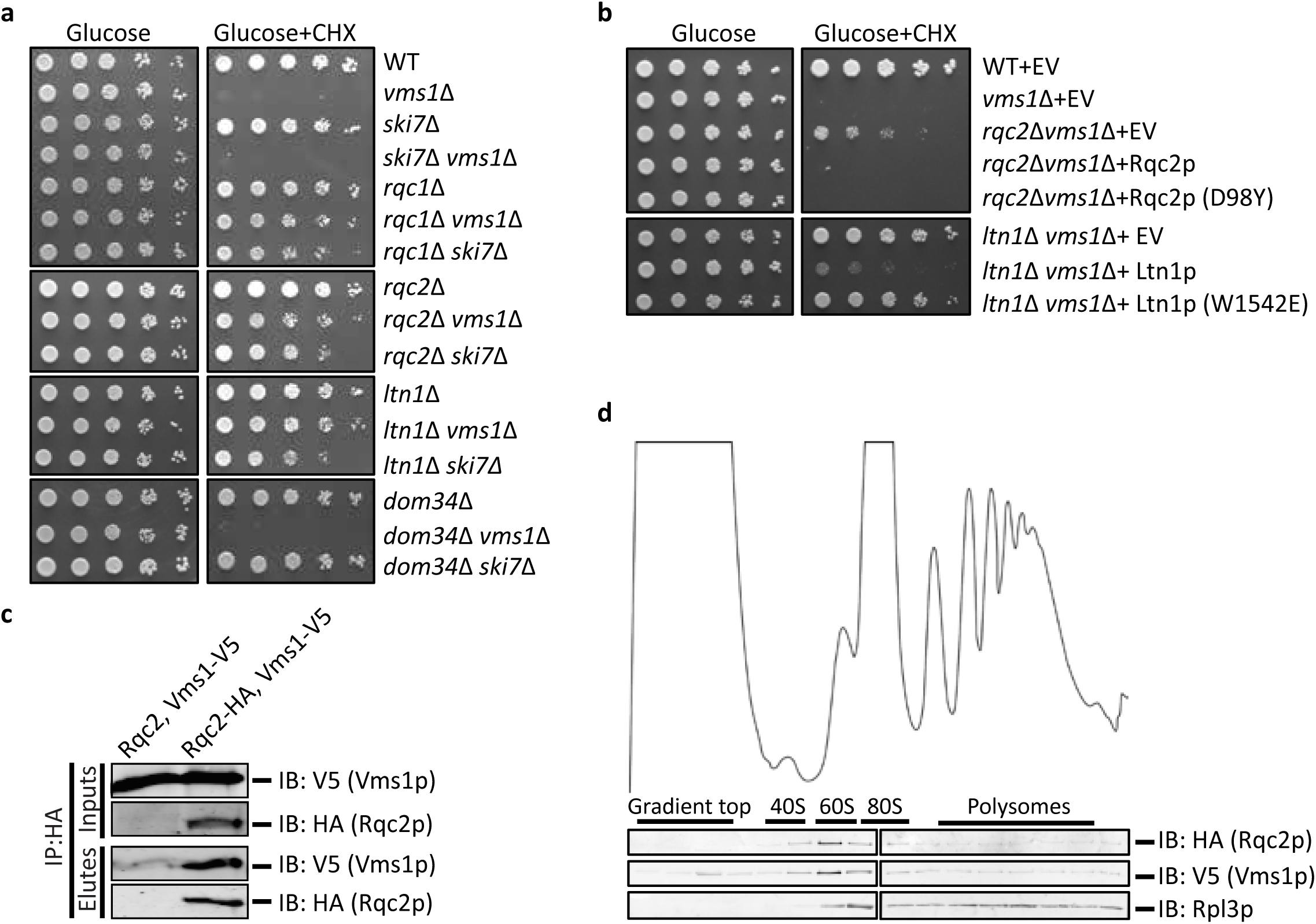
Vms1 physically and genetically interacts with the RQC. (a, b) Serial dilutions of indicated strains were spotted on media containing glucose or glucose supplemented with cycloheximide (CHX). EV, empty vector. (c) Immunoprecipitation using anti-HA antibody in the strains *rqc2*Δ *vms1*Δ expressing Rqc2p and Vms1p-V5 (control) or Rqc2p-HA and Vms1p-V5. Immunoblotting of HA and V5 were used to identify Rqc2p and Vms1p, respectively. (d) Polysome profile of the *rqc2*Δ *vms1*Δ strain expressing Rqc2p-HA and Vms1p-V5 treated with CHX prior to fractionation using sucrose density centrifugation. The sedimentation of ribosomal particles was inferred from the *A*_254_ profile (40S, 60S, 80S and polysomes) and the distribution of the 60S subunit was confirmed by immunoblotting of the ribosomal subunit, Rpl3p. Immunoblotting of HA and V5 was used to detect Rqc2p and Vms1p, respectively.

The RQC assembles on a failed 60S subunit to both ubiquitinate and elongate nascent polypeptides with CAT tails. To determine whether one or both of these activities generate the putative toxic RQC product, we eliminated them separately by expressing a CAT-tailing-defective Rqc2p mutant (*RQC2^D98Y^*)^4,8,14^ or a variant of Ltn1p deficient in ubiquitination (*LTN1*^*W1542E*^)^10,21^. CAT-tailing by Rqc2p was dispensable, but ubiquitination by Ltn1p was essential, for conferring CHX sensitivity on a *vms1*Δ mutant strain (**Fig. 1b**). Similar to *RQC2* and *LTN1* (**Fig. 1b**), plasmid expression of wild type *RQC1* was sufficient to reverse the rescue of *vms1*Δ cycloheximide sensitivity conferred by *RQC1* deletion (**Extended Data Fig. 1a**).

In light of the association between Vms1 activity and mitochondrial stress, we examined the ability of these single and double mutant strains to grow in glycerol medium, which requires mitochondrial respiration. Interestingly, deletion of *LTN1*—but not *RQC1*, *RQC2* or *DOM34*—strongly impaired glycerol growth of *vms1*Δ cells (**Extended Data Fig. 1b**). Similarly, *ski7*Δ *vms1*Δ double mutant cells also exhibited partially impaired glycerol growth (**Extended Data Fig. 1b**). These data indicate a specific relationship between Vms1p, RQC function, and mitochondrial homeostasis, which is consistent with a recent report that stalled polypeptides that cannot be ubiquitinated by Ltn1p accumulate within mitochondria^22^.

These genetic interactions prompted us to determine whether Vms1p physically interacts with members of the RQC, as has been reported previously^1,3,22^. As expected, isolation of Rqc2p-HA co-immunoprecipitated Vms1p-V5 whereas Rqc1p-HA or Ltn1p-HA showed minimal Vms1p interaction (**Fig. 1c and Extended Data Fig. 1c-d**). Consistently, both Rqc2p and Vms1p co-migrated with the 60S ribosome subunit during sucrose gradient sedimentation following CHX treatment (**Fig. 1d, Extended Data Fig. 1e**). Co-migration of Vms1p with the 60S ribosome was not affected by either deletion of *RQC2* or by expression of WT or a CAT-tailing defective, D98Y, mutant of Rqc2p (**Extended Data Fig. 1e**). Similarly, neither deletion nor overexpression of Vms1p from the strong *GAL1* promoter in galactose medium had any effect on the co-migration of Rqc2p with the 60S ribosomal subunit (**Extended Data Fig. 1e,f**).

These genetic and physical interactions motivated us to evaluate whether RQC substrates accumulate in *VMS1* mutant cells. We utilized a well-characterized mRNA that encodes FLAG-tagged GFP followed by a hammerhead ribozyme that self-cleaves in vivo (FLAG-GFP^Rz^, **Fig. 2a**)^4,23^ to generate a truncated mRNA encoding FLAG-GFP without a stop codon or poly-A tail. Translation of this mRNA triggers ribosome stalling and targeting to the RQC system. Deletion of *SKI7* enhances expression of GFP by inhibiting degradation of the cleaved mRNA^24^. We confirmed that deletion of *RQC1*, *RQC2*, and *LTN1* each lead to accumulation of FLAG-GFP^Rz^, whereas the nascent chain failed to accumulate in the *ski7*Δ single mutant (**Fig. 2a-c, Extended Data Fig. 2a**). Loss of Vms1p also led to accumulation of FLAG-GFP^Rz^ to a level similar to that observed for the core RQC components (**Fig. 2a-c, Extended Data Fig. 2b)**. Combination of *VMS1* deletion with the deletion of *RQC1*, *RQC2* and *LTN1* had no additive effect on GFP accumulation (**Fig. 2b**). Immunoblot analysis showed similar results for the single, double and triple mutant strains, in which RQC2-dependent, high molecular weight aggregates are also apparent (**Fig. 2c, Extended Data Fig. 2b**)^12–14^. Loss of *DOM34* led to decreased accumulation of GFP fluorescence, even in the *vms1*Δ strain, consistent with Dom34p’s upstream role in ribosome splitting and suggestive of alternative pathways for degrading nascent chains when the Dom34p/Hbs1p subunit splitting activity is unavailable (**Fig. 2a-c**). Interestingly, accumulation of FLAG-GFP^Rz^ in *vms1*Δ mutant cells occurs despite lower mRNA abundance (**Extended Data Fig. 2c**).

**Figure 2.**
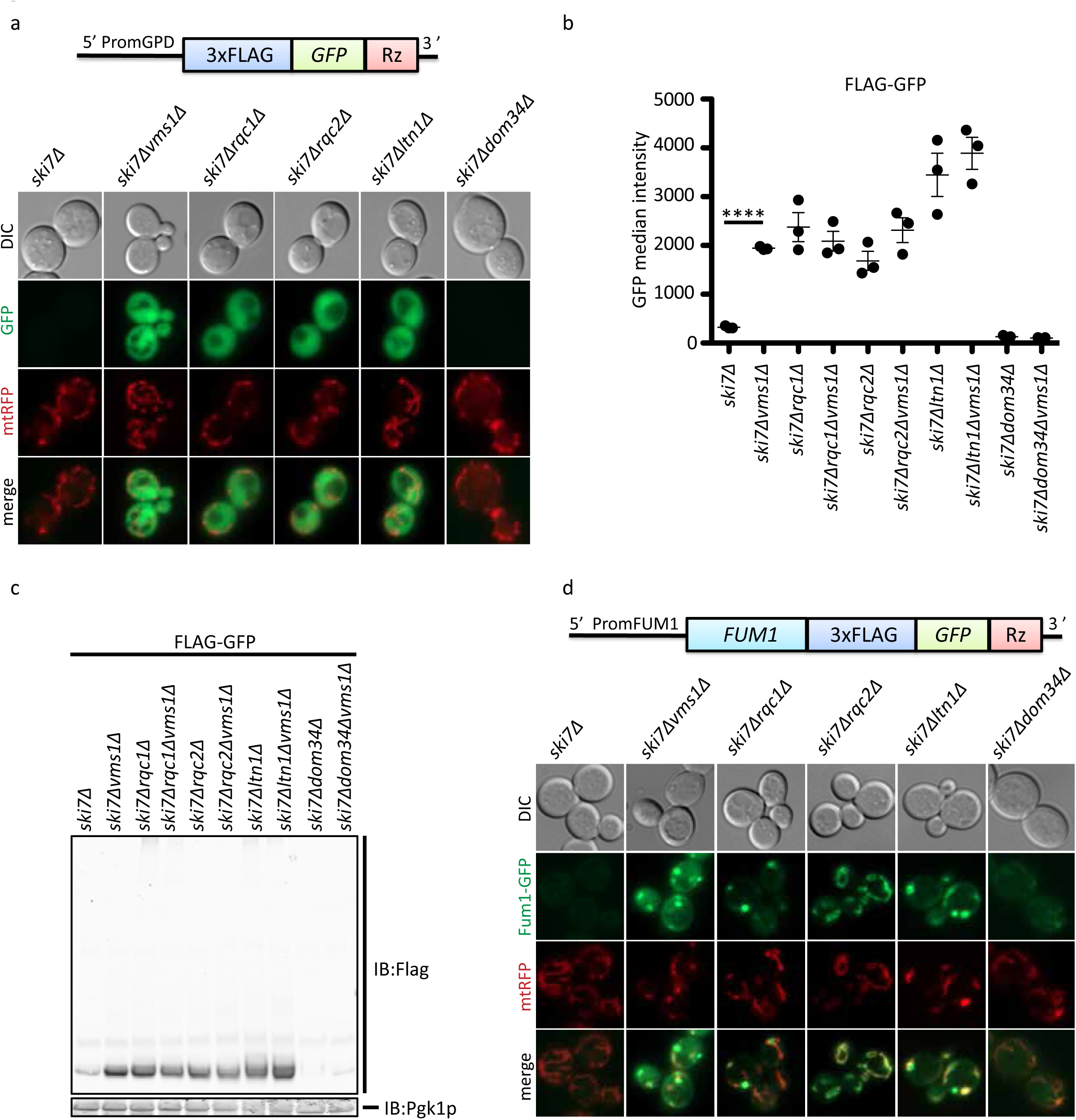
Vms1 is required for resolving RQC substrates. (a) Fluorescence microscopy analysis of the indicated strains expressing the FLAG-GFP^Rz^ construct under the GPD promoter and the mitochondrial marker, mtRFP. (b) Flow cytometry quantifications of FLAG-GFP accumulation in the indicated strains. Median GFP intensity values are plotted (*n*=3, mean ± s.e.m. ^⋆⋆⋆⋆^ *P* < 0.0001, *P*-value was calculated using unpaired Student’s t-test). (c) Immunoblot analysis of indicated strains expressing the FLAG-GFP^Rz^ construct. Immunoblotting of Flag was used to detect the accumulation of the stalled construct. Pgklp was used as loading control. (d) Fluorescence microscopy analysis of the indicated strains expressing the Fum1-FLAG-GFP^Rz^ construct expressed from the *FUM1* endogenous promoter and the mitochondrial marker, mtRFP.

In addition to the FLAG-GFP^Rz^ construct, which generates a cytosolic RQC substrate, we also examined RQC activity on fumarase, which is encoded by the *FUM1* gene and co-translationally imported into the mitochondria^25^. As with FLAG-GFP^Rz^, fluorescence from the Fum1-FLAG-GFP^Rz^ construct, expressed from the native *FUM1* promoter, was also maintained at a low level in the *ski7*Δ mutant strain (**Fig. 2d**). Deletion of *VMS1*, *RQC1*, *RQC2*, or *LTN1* each led to profound accumulation of GFP fluorescence, almost all of which colocalized with mitochondria-targeted RFP (mtRFP). The *vms1*Δ, *rqc1*Δ and *ltn1*Δ mutants, which retain Rqc2p and CAT-tailing activity, all exhibited Fum1-GFP aggregates within or near mitochondria, comparable to recent observations of other mitochondria-destined nascent chains^22^ (**Fig. 2d**). The *rqc2*Δ mutant exhibited a more uniformly mitochondrial localization pattern (**Fig. 2d**), consistent with the model that CAT-tailing mediates intra-mitochondrial aggregation of polypeptides that stall during co-translational import^22^. Together, these data demonstrate that Vms1p is required for the degradation of substrates derived from truncated mRNAs, whether they are destined for an organelle or the cytosol.

Understanding of how Vms1p facilitates the clearance of stalled translation products was guided by our recent crystal structure determination of *S. cerevisiae* Vms1p (**Fig. 3a-b**)^26^ This structure includes the highly conserved central region of Vms1p, which we named the Mitochondrial Targeting Domain (MTD) because it is necessary and sufficient for mitochondrial localization in response to stress^15^. This localization activity requires a hydrophobic groove along the bottom of the MTD and direct binding to ergosterol peroxide^26^. Intriguingly, the Vms1p MTD structure resembles structures of the catalytic domain of eukaryotic peptide chain release factor subunit 1 (eRF1), as well as Dom34p and RNaseE, which both resemble tRNA hydrolases^27–30^ (**Fig. 3b**, **Extended Data Fig. 3a**). The only region of the Vms1p MTD that diverges substantially from the release factor fold is the face of the MTD that mediates mitochondrial localization^26^ (**Fig. 3b**). The loop of eRF1 that harbors the signature GGQ motif—which catalyzes the hydrolytic attack on the peptidyl-tRNA ester bond—and the orthologous loop of the Vms1p MTD, can both be unstructured when not bound to the ribosome^28,30^ (**Fig. 3b**).

**Figure 3.**
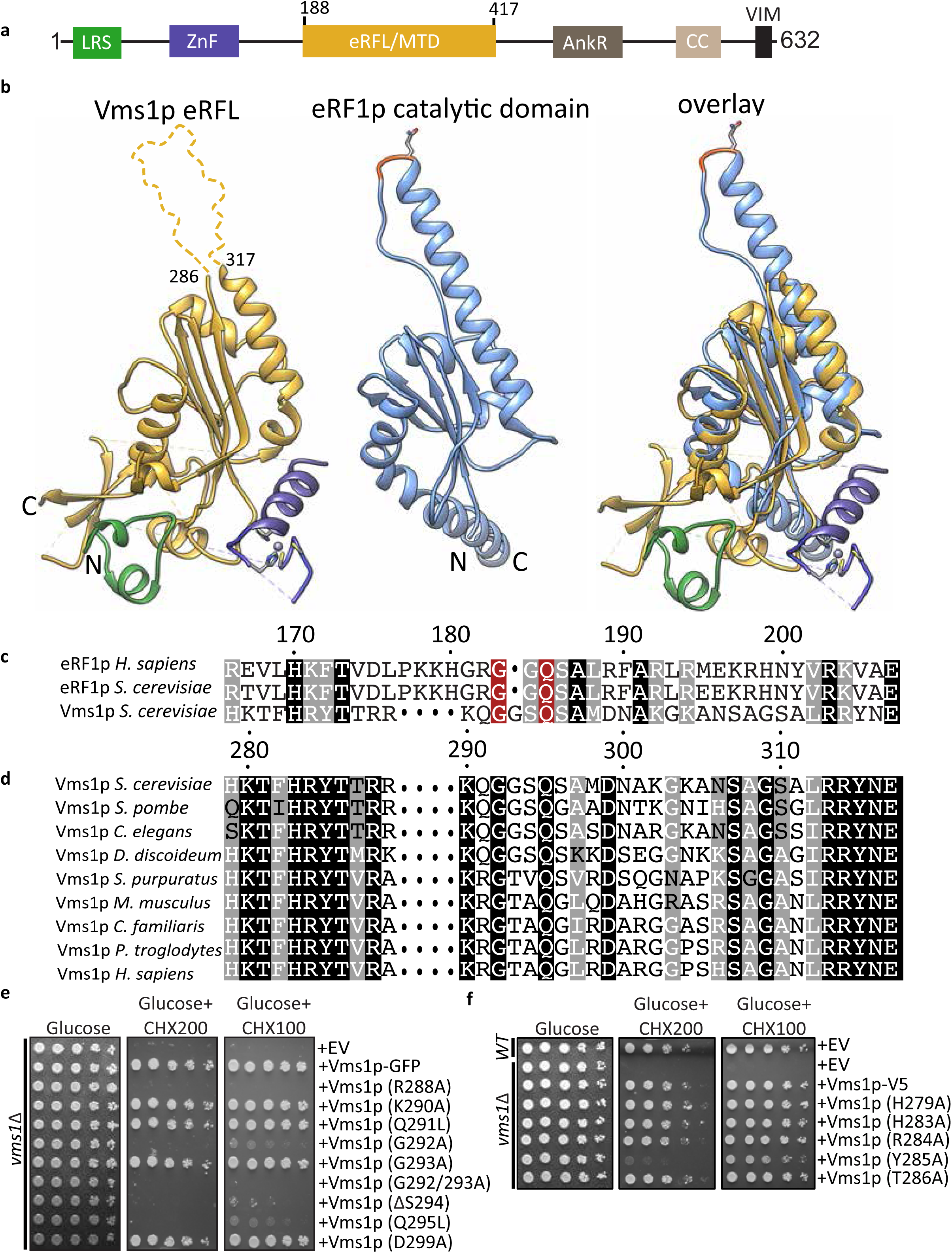
Vms1 is structurally homologous to tRNA hydrolases. (a) Domain structure of Vms1p. LRS, Leucine Rich Sequence; ZnF, Zinc Finger; MTD/eRFL, Mitochondrial Targeting Domain/eRF1-like; AnkR, Ankryin Repeat; CC, Coil-Coil; VIM, VCP-Interacting Motif. Residues 188-417 represent the MTD/eRFL boundaries. (b) Structural alignment of Vms1p (left, 5WHG) and eRF1p (middle, 3JAHii, residues 144280). Dashed lines indicate connections made by residues that are not resolved in the Vms1p crystal structure. The GGQ (red) loop of eRF1p is ordered in the ribosome-bound structure shown here. (c) Sequence alignment of Vms1p and eRF1p. White letters with gray, black, or red background indicates similarity, identity, or GxxQ residues, respectively. (d) Sequence alignment of Vms1p orthologs across the GxxQ region. Coloring as in (c). (e,f) Serial dilutions of indicated strains were spotted on media containing glucose or glucose supplemented with cycloheximide (CHX). EV, empty vector.

Sequence alignment showed that although Vms1p lacks a strict GGQ motif characteristic of eRF1p, it does possess an invariant glutamine that aligns with the catalytic glutamine of eRF1 (**Fig. 3c**). In yeast Vms1p, that glutamine residue is embedded within a GGSQ motif that is reminiscent of the eRF1 catalytic GGQ, while in other species the conservation other than the initial glycine and glutamine is less apparent (**Fig. 3c-d**). The Vms1p MTD lacks similarity to the non-catalytic eRF1 domain 1, which discriminates stop codons from sense codons^30^. This is consistent with Vms1p functioning in stop codon-independent tRNA hydrolysis within a 60S, rather than 80S, ribosome.

These observations inspired us to determine whether Vms1p enables the extraction of failed translation products from the stalled 60S by hydrolyzing the ester bond anchoring them to tRNA. We first asked which residues and regions are required for the genetic functions of *VMS1.* We first tested an HbϕT motif just N-terminal to the conserved ‘GxxQ’ motif that mediates ribosome interactions of eRF1 (**Fig. 3c-d**). Vms1p mutants of these residues, H279A, H283A, R284A and T286A, were indistinguishable from WT, while the Y285A mutant exhibited a partial CHX sensitivity (**Fig. 3f**). In contrast, mutation of the ‘GxxQ’ residues G292 and Q295 and the highly conserved R288 residue conferred strong loss-of-function phenotypes (**Fig. 3e**). Deletion of S294 to convert the GGSQ of *S. cerevisiae* Vms1p into a GGQ motif, as in eRF1, abrogated *VMS1* function (**Figure 3e**). While all of these ‘GxxQ’ ‘mutants failed to confer resistance to 200 ng/ml CHX, only the R288A and G292A/G293A mutants were inactive at the lowest (100 ng/ml) concentration of CHX tested (**Figure 3e**). Interestingly, both of these mutants also failed to rescue glycerol growth in an *ltn1*Δ *vms1*Δ double mutant, whereas wild-type *VMS1* and the other mutants did rescue growth (**Extended Data Fig. 3b**). The R288A, GG292/3AA and Q295L mutants also exhibited enhanced accumulation of FLAG-GFP^Rz^ in the *ski7*Δ background similar to the *vms1*Δ mutant (**Extended Data Fig. 3d**). Importantly, the Vms1p mutants interact normally, if not more strongly, with Rqc2p based on co-immunoprecipitation experiments (**Extended Data Fig. 3e**). In light of these observations, we hereafter refer to the MTD as the MTD/eRFL domain, where eRFL refers to eRF1-like.

In addition to these loop residues, the ability of Vms1p to confer complete CHX resistance in both *vms1*Δ and *ski7*Δ *vms1*Δ also required the VCP-interacting motif (VIM), which mediates interaction with Cdc48p/VCP/p97^16^ (**Extended Data Fig. 3c**). Interestingly, the VIM is not required for growth of the *ltn1*Δ *vms1*Δ double mutant on glycerol (**Extended Data Fig. 3c**), which indicates that mitochondrial homeostasis can be maintained even without Cdc48p binding.

To directly test whether Vms1p catalyzes peptidyl-tRNA hydrolysis, we utilized our recently described *S. cerevisiae* in vitro translation (ScIVT) system to monitor the synthesis and fate of a robust stalling reporter and its peptidyl-tRNA intermediate^10^. RQC-intact extracts translate this reporter, split the stalled 80S ribosome into constituent 60S and 40S subunits, elongate the nascent chain with a CAT tail, and ubiquitinate exposed Lys residues. These extracts also hydrolyze the peptidyl-tRNA ester bond to generate the released polypeptide (**Fig. 4a**)^10^. We observed that extracts prepared from *vms1*Δ mutant cells also produced peptidyl-tRNA conjugates, but loss of the peptidyl-tRNA species and appearance of the released translation product were slower than in WT extracts (**Fig. 4a-b**). This is somewhat obscured by the fact that the *vms1*Δ mutant has lower overall translation, which leads to a decreased amount of the free nascent chain and peptidyl-tRNA conjugates. We performed a similar experiment in the *ski7*Δ and *ski7*Δ *vms1*Δ mutant strains and found that in the *ski7*Δ background the deletion of *VMS1* conferred a much more obvious stabilization of the peptidyl-tRNA species and qualitatively delayed release of the polypeptides (**Fig. 4c-d**). In this *ski7*Δ background, deletion of *RQC2* conferred a modest stabilization of the peptidyl-tRNA conjugate^10^ and deleting *RQC2* had little effect on the *ski7*Δ *vms1*Δ double mutant (**Extended Data Fig. 4a**). We next purified full-length and C-terminally truncated (1-417) *S. cerevisiae* Vms1p and found that each of these proteins dramatically accelerated the production of the released polypeptide in a dose-dependent manner in WT, *rqc2*Δ and *vms1*Δ extracts (**Fig. 4e, Extended Data Fig. 4b**). Importantly, the 1-417 truncation mutant lacks the C-terminal VIM domain and is unable to interact with Cdc48p (**Extended data Fig. 3e**). We therefore conclude that while the Vms1-Cdc48 interaction is important for CHX resistance and other RQC-related functions, it is dispensable for peptidyl-tRNA hydrolysis.

**Figure 4.**
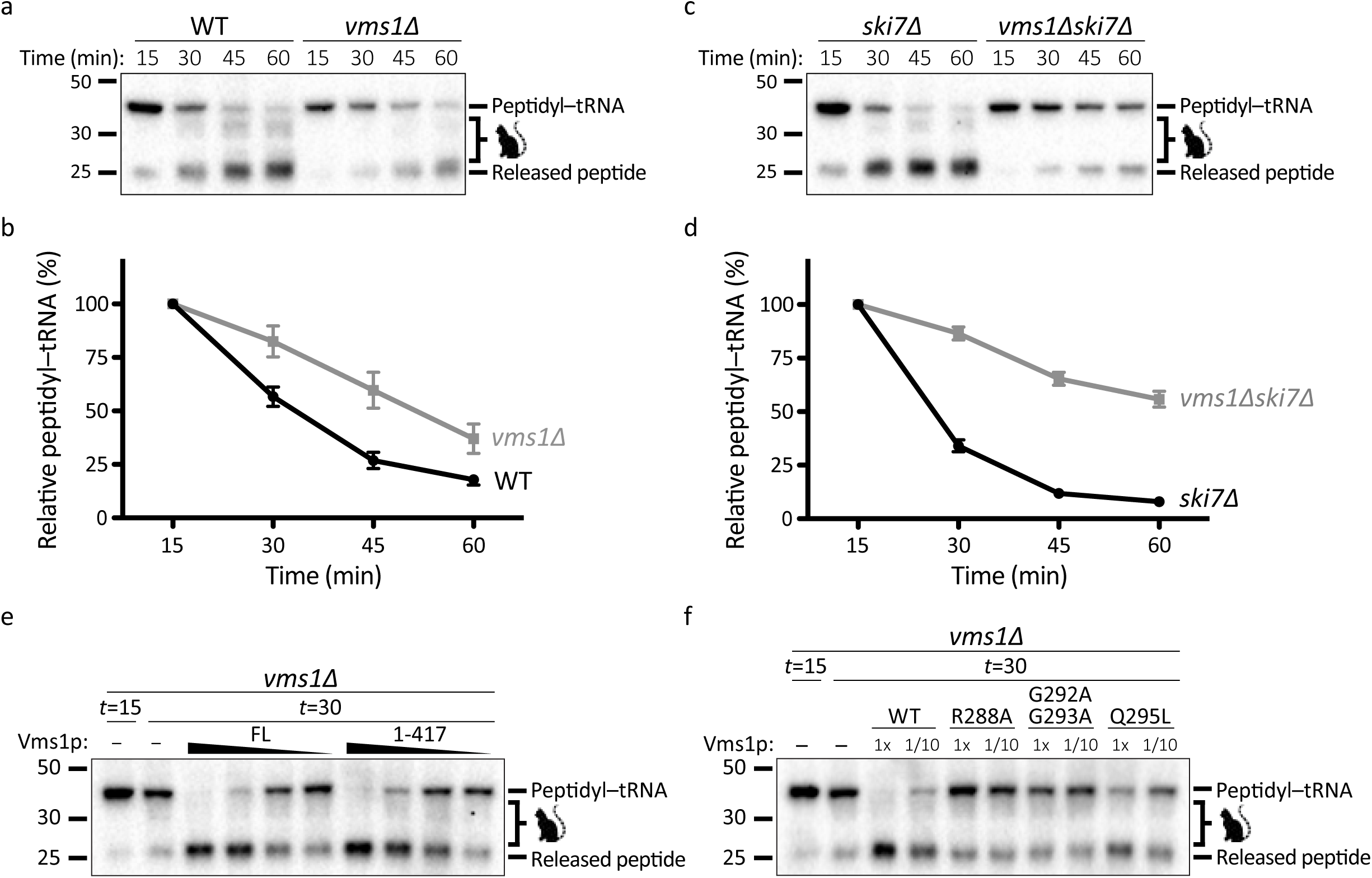
Vms1p exhibits tRNA hydrolase activity towards RQC substrates. (a) Time courses of *S. cerevisiae* in vitro translation (ScIVT) reactions prepared with a truncated mRNA (lacking a stop codon). Extract genotypes are indicated above. Peptides that have been CAT-tailed and released are denoted by: 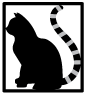 (b) Quantification of peptidyl-tRNA species in (a). Mean ± s.e.m, *n*=6. ^⋆⋆⋆⋆^ *P* < 0.0001. *P*-value was calculated using a 2-way ANOVA. (c) Time courses of ScIVT reactions prepared as in (a). (d) Quantification of peptidyl-tRNA species in (c). Mean ± s.e.m, *n*=8. ^⋆⋆⋆⋆^ *P* < 0.0001. *P*-value was calculated using a 2-way ANOVA. (e) ScIVT reactions prepared as in (a) with a *vms1Δ* extract. At *t* = 5, buffer (-) or pure protein was added. Slopes indicate a titration series of decreasing protein concentrations (see Methods). FL = Full Length Vms1; 1-417 = N-terminus through eRFL domain. (f) ScIVT reactions prepared as in (a) with a *vms1Δ* extract. At *t* = 15, buffer, WT(1-417) protein, or mutant(1-417) protein was added.

We next tested release factor activity of the MTD/eRFL domain structure-based mutants described above. R288A and G292A/G293A mutants had no hydrolysis activity, even at 10-fold higher concentration than the concentration at which WT Vms1p catalyzed complete tRNA release (**Fig. 4f**). The Q295L, G292A, and ΔS294 mutants also exhibited strongly impaired release factor activity (**Fig. 4f and Extended Data Fig. 4c**).

Our data have identified a key constituent of the RQC pathway in Eukarya: a tRNA hydrolase that liberates failed polypeptides from the aberrant 60S:peptidyl-tRNA species that accumulate when ribosomes stall and split apart. Without this activity, translation products remain anchored in 60S ribosomes, which therefore cannot be recycled for future use. The dual functions of the Vms1p MTD/eRFL identified here as an RQC pathway release factor and earlier as a targeting domain in mitochondrial stress responses, portend exciting future work at the intersection of proteostasis and organelle homeostasis. We have previously reported that Vms1p localizes to mitochondria under conditions of mitochondrial damage or cellular stressors, including rapamycin treatment, by binding to the oxidized sterol ergosterol peroxide^15,16,26^. The MTD/eRFL domain is necessary and sufficient for this localization, which is mediated by a direct interaction between a face of the MTD/eRFL domain that should remain exposed even when the domain is in the ‘A-site’ of the 60S and the catalytic GGSQ loop is presumably reaching into the peptidyl transferase center to catalyze hydrolysis of the peptidyl-tRNA ester bond. While the relationship between mitochondrial localization and RQC-coupled release factor activity remains unclear, it is intriguing to speculate that this is indicative of a role for Vms1p—and the RQC as a whole—in the response to mitochondrial stress. Consistent with this possibility, *ltn1*Δ *vms1*Δ and *ski7*Δ *vms1*Δ double mutant cells exhibit impaired glycerol growth, which correlates with impaired mitochondrial respiration^22^. We therefore propose that Vms1p is required for the resolution of peptidyl-tRNA conjugates of stalled nascent chains in the cytosol as well as those destined for organelles like the mitochondria, where it might mediate a particular role in protecting mitochondria from proteostasis challenges.

## Methods

### Yeast strains and growth conditions

*Saccharomyces cerevisiae* BY4741 (*MATa*, *his3 leu2 met15 ura3*) was used as the wild-type strain. Each mutant was generated in diploid cells using a standard PCR-based homologous recombination method. The genotypes of all strains used in this study are listed in **Extended Data Table 1A.** Yeast transformations were performed by the standard TE/LiAc method and transformed cells were recovered and grown in synthetic complete glucose (SD) medium lacking the appropriate amino acid(s) for selection. The medium used included YPA and synthetic minimal medium supplemented with 2% glucose or 3% glycerol. Cycloheximide was added at a final concentration of 100 ng/ml or 200 ng/ml when indicated.

All plasmid constructs were generated by PCR and cloned into the yeast expression vectors pRS413, pRS14 or pRS416 as indicated in **Extended Data Table 1B**.

Growth assays were performed using synthetic minimal media supplemented with the appropriate amino acids and indicated carbon source. For plate-based growth assays, overnight cultures were back-diluted to equivalent ODs and spotted as 10-fold serial dilutions. Cells were grown at 30°C.

### Immunoprecipitations

p^VMs1^-VMS1-V5 (or VMS1 mutant) was co-expressed with an endogenous promoter-HIS6-HA2 tagged RQC component (RQC1, RQC2, LTN1) in the cognate double mutant strain. ~50 OD were harvested in log phase and resuspended in lP buffer (20 mM Tris pH 7.4, 50 mM NaCl, 0.2% TritonX-100), vortexed 10 X 1min, clarified via centrifugation, and added to anti-HA magnetic beads (Thermo scientific #88836). After 4 hours of incubation, beads were pelleted via magnet and washed 4X with 1 ml of IP buffer. Proteins were eluted with 50 μl of 2X Laemmli buffer (20% glycerol, 125 mM Tris-HCl pH 6.8, 4% SDS, 0.02% bromophenol blue).

### Polysome profiling

Yeast cultures were grown to OD_600_ ~1, cycloheximide was added to a final concentration of 0.05mg/ml, and cells were harvested by centrifugation five minutes later. Cell pellets were washed in buffer A (20 mM Tris-HCl pH 7.4, 50 mM KCl, 10 mM MgCl_2_, 1 mM DTT, 100 μg/mL cycloheximide, 1X RNAsecure [Ambion], and 1X yeast protease inhibitor [Sigma]). Pellets were weighted and resuspended in 1.3 volumes of Buffer A. An equal volume of glass beads was added and suspensions were vortexed for 30 secs a total of 8 times interspersed with 1 min. incubation on ice. Following centrifugation at 3,000 x g for 5 min, supernatant was centrifuged at 11,300 x g for 2 min at 4°C, after which supernatant was centrifuged at 11,300 x g for 10 min. Protein extracts were overlaid onto a linear sucrose gradient of 15-50% and centrifuged at 234,600 x g for 90 min. The gradients were passed through a continuous-flow chamber and monitored at 254 nm with a UV absorbance detector (ISCO UA-6) to obtain ribosomal profiles. Fractions (16) were collected, resuspended in 2X Laemmli sample buffer supplemented with 2.5% beta-mercaptoethanol, and analyzed by western blotting.

### SDS-PAGE

Whole Cell Extracts were prepared from 3-5 OD of cells at OD_600_~ 1.5 by solubilization in 250 μl of 2 M LiAc, incubated for 8 min on ice followed by centrifugation at 0.9 x g for 5 minutes at 4°C. The pellet was resuspended in 250 μl of 0.4 M NaOH and incubated on ice for 8 min. followed by centrifugation at 16,000 x g for 3 min. Next, the pellet was resuspended in 1X Laemmli buffer with 2.5% beta-mercaptoethanol, boiled for 5 minutes, and centrifuged at 0.9 x g for 1 min. Supernatants were collected and loaded onto acrylamide:bisacrylamide (37.5:1) gels. Subsequent immunoblotting was done with the indicated antibodies: HA (PRB-101C-200), V5 (ab9116), FLAG (F7425), Pgk1: (ab113687) and Rpl3 (scRPL3).

### Fluorescence microscopy

WT (BY4741) or derived mutant strains were transformed with a plasmid expressing mitochondria-targeted (ATPase subunit, Su9) RFP, mtRFP, and plasmids expressing 6XFlag3XHis-GFP-Rz or FUM1-6XFlag3XHis-GFP-Rz under the *GPD* or native *FUM1* promoter, respectively. The cells were grown to mid-log phase and imaged using the Axio Observer Z1 imaging system (Carl Zeiss). Digital fluorescence and differential interference contrast (DIC) images were acquired using a monochrome digital camera (AxioCam MRm) and analyzed using the Zen 2 software (Carl Zeiss).

### Fluorescence assisted cell sorting

GFP-expressing strains and untransformed control were grown to OD_600_~ 1 and pelleted by centrifugation at 100 x g for 5 min. Cell pellets were washed once in 1X PBS buffer, resuspended in 1 ml of 1X PBS, and analyzed using the BDFACSCanto Analyzer (488 laser and optical filter FITC). 30,000 events were measured and the median values of three independent biological replicates were analyzed by unpaired *Student t*-test (two-tailed) confidence interval value set to p<0.05. Error bars represent standard error of the mean. This analysis was done using the statistics software: Graphpad Prism 6.

### Protein Expression and Purification

For the His_12_ tagged proteins, constructs were transformed into JRY1734 (pep4::HIS3 prb1::LEU2 bar1::HISG lys2::GAL1/10-GAL4) and grown in synthetic media lacking Uracil with 3% glycerol and 2% ethanol. When the OD_600_ reached ~0.5, 0.5% galactose was added to the cultures, which were grown for another 6 hours before harvesting by centrifugation, washing of the pellet with sterile H2O, and flash freezing in liquid nitrogen. Cells were lysed using a pulverizer (SPEX SamplePrep 6870), and the lysed powder was thoroughly resuspended in lysis buffer (20 mM Tris pH 8.0, 300 mM NaCl, 5% glycerol) supplemented with protease inhibitors (aprotinin, leupeptin, pepstatin A, and PMSF) (Sigma). The resuspended lysate was clarified by centrifugation and added to Ni-NTA resin (Qiagen #30250) for 1 hour, washed with 10 CV of lysis buffer, 10 CV of lysis buffer with 40 mM imidazole, and eluted with lysis buffer made up with 250 mM imidazole. Eluted protein was dialyzed into IVT-compatible buffer (20 mM HEPES-KOH pH7.4, 150mM KOAc, 5% glycerol, 2mM DTT) and concentrated.

### *Saccharomyces cerevisiae* in vitro translation (ScIVT)

Preparation of in vitro translation extracts, mRNA, and in vitro translation reactions was performed as previously described^10^. Briefly, *S. cerevisiae* strains were cryo-lysed and cell debris was cleared by sequential centrifugation before dialysis into fresh lysis buffer. mRNAs were generated by run-off transcription from PCR-amplified templates of 3xHA-NanoLuciferase to produce transcripts lacking a stop codon and 3’UTR (truncated quality control substrate). Transcription products were capped and extracted prior to freezing for use in ScIVTs. For ScIVT reactions, extracts were first treated with MNase to remove endogenous mRNAs and then supplemented with 480 ng mRNA to initiate translation. Reaction aliquots were sampled at indicated time points by quenching in 2X Laemmli Sample Buffer. Proteins were separated by SDS-PAGE, and HA-tagged translation products were visualized by immunoblotting (Roche3F10). To quantify release, the abundance of peptidyl-tRNA was measured with Fiji (https://imagei.net/Fiji) and normalized as percentage of the initial 15 min time point. Mean values of at least 2 biological replicates and 2-3 technical replicates were analyzed and plotted in Prism (GraphPad software). *P*-values were calculated using a 2-way ANOVA as follows: WT vs. *vms1*Δ, *F*= 28.62, *DFn*= 1 and *F*= 76.42, *DFn*= 3 for genotype and time, respectively, and *ski7*Δ vs. *ski7*Δ *vms1*Δ, *F*= 561.69, *DFn*= 1 and *F*= 357.63, *DFn*= 3 for genotype and time, respectively. Error bars represent standard error of the mean. The ramps of panel (e) represent a decreasing titration series of 4.2 μM, 0.42 μM, 0.21 μM and 0.105 μM final protein concentrations. In Figure 4f and Extended Data Figure 4c, 1x and 1/10 refer to final protein concentrations of 4.2 μM and 0.42 μM, respectively.

### Quantitative RT-PCR

RNA was purified from 40 ml of yeast cultures grown to OD_600_~ 1. Pelleted cells were washed once with water and resuspended in 700 μl of Trizol reagent (Ambion). An equal volume of glass beads was added and suspensions were vortexed for 30 secs intervened with 1 min. rest intervals. Next, the manufacture’s protocol from the Direct-zol kit (Zymo research: R2050-11-330) was followed. cDNA was obtained from 0.5 μg of purified RNA using the High-capacity cDNA Reverse Transcription kit (4368814) from Applied Biosystems. qPCR was performed using the LightCycler 480 SYBR Green I Master (04707516001) from Roche and using a FLAG-HIS and Actin primer pairs. qPCR analysis was done by Absolute Quantification/2^nd^ derivative of three independent biological replicates, each performed in triplicate. The statistical analysis of mRNA transcript abundance was done after normalization with Actin. The statistics software Graphpad Prism 6 was used to performed a *Student* t-test (unpaired two-tail) with a confidence interval value of p<0.05. Error bars represent standard error of the mean.

## Data availability

The authors declare that the data supporting the findings of this study are available within the paper and its supplementary information files.

Further relevant data on the genes studied in this manuscript (*VMS1:* YDR049W, *RQC1:* YDR333C, *RQC2:* YPL009C, *LTN1:* YMR2476, *SKI7:* YOR076C, *DOM34:* YNL01W) can be found at: **https://www.yeastgenome.org**

## Acknowledgements

This work was supported by a Faculty Scholar grant from the Howard Hughes Medical Institute (A.F.), the Searle Scholars Program (A.F.), NIH grant GM115129 (to C.P.H. and J.R.), a grant from the Nora Eccles Treadwell Foundation (to J.R. and C.P.H.), NIH grant 1DP2GM110772-01 (A.F.), training grants T32HL007576, AHA 14POST20380216, and T32DK007115 (E.K.F.), a Hillblom Graduate Research fellowship (C.J.H.), a Heyman Discovery fellowship (B.A.O), and the Howard Hughes Medical Institute (J.R.). A.F. is a Chan Zuckerberg Biohub investigator. This work was supported by the University of Utah Flow Cytometry Facility in addition to the National Cancer Institute through Award Number 5P30CA042014-24.

## Author contributions

O.Z.R., E.K.F., C.J.H., J.R, and A.F. designed the study and wrote the manuscript. N.D.T. and J.V.V. ran the polysome assays. O.Z.R., E.K.F., C.J.H. collected the data. J.V.V. and S.F. generated plasmid constructs and yeast strains. C.P.H helped determine and analyze the Vms1 structure. R.K. performed structural homology modeling and alignments. B.A.O helped with the IVT assays. P.S. helped with the co-immunoprecipitation experiments. All authors commented and approved of the final manuscript.

## Author information

Authors declare no competing interests.

---

**Extended Data Figure 1.**
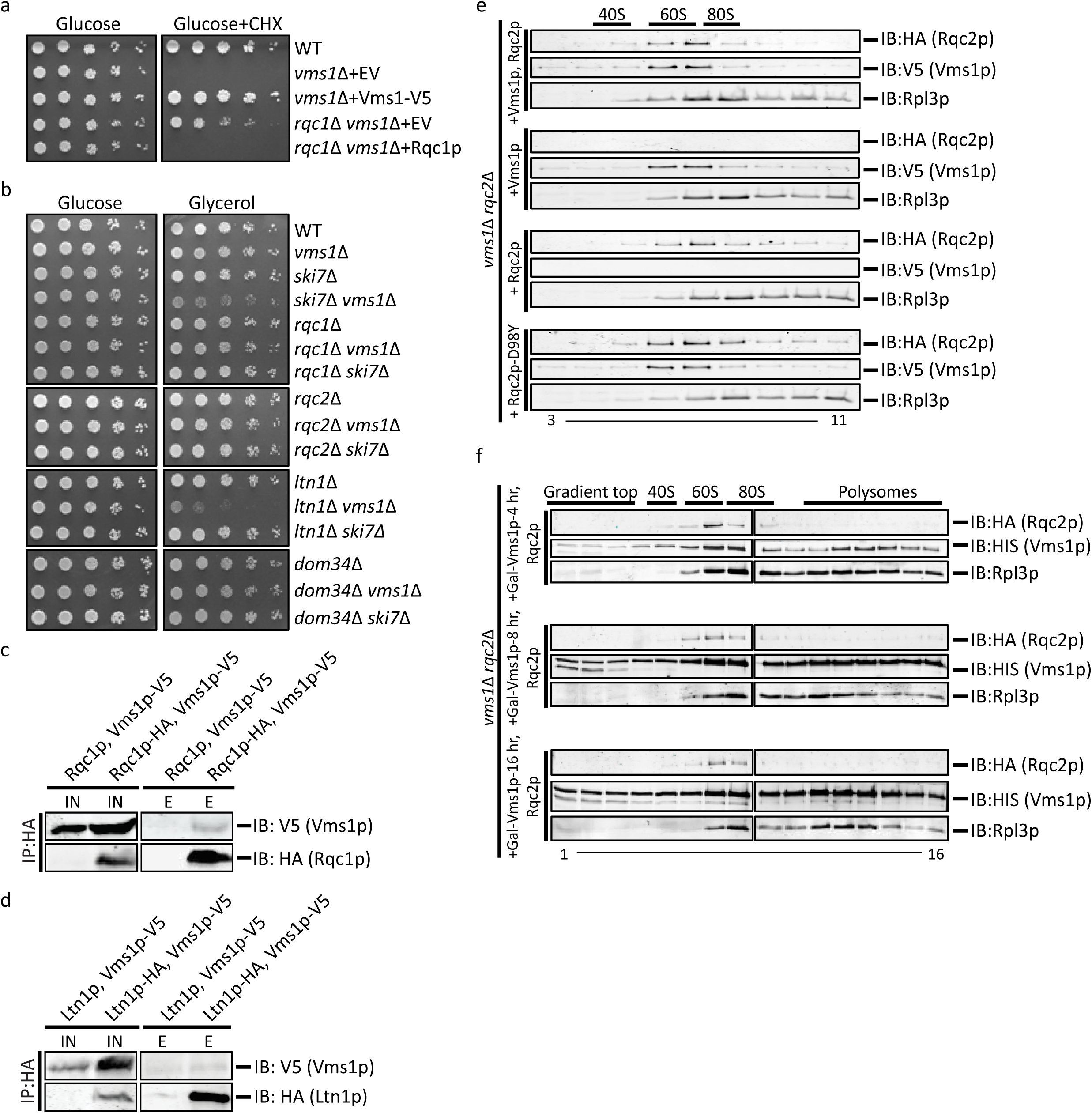
(a) Serial dilutions of indicated strains were spotted on media containing glucose or glucose supplemented with cycloheximide (CHX) and grown for 2 or 3 days, respectively. (b) Serial dilutions of the indicated strains were spotted on medium containing glucose or glycerol and grown for 2 or 3 days, respectively. (c) Immunoprecipitations using anti-HA antibody in the *rqc1*Δ *vms1*Δ strain expressing Rqc1p and Vms1p-V5 (control) or Rqc1p-HA and Vms1p-V5. Immunoblotting of HA and V5 were used to identify Rqc1p and Vms1p, respectively. (d) Immunoprecipitations using anti-HA antibody in the *ltn1*Δ *vms1*Δ strain expressing Ltn1p and Vms1p-V5 (control) or Ltn1p-HA and Vms1p-V5. Immunoblotting of HA and V5 were used to identify Ltn1p and Vms1p, respectively. (e) Polysome profiles of whole cell extracts from the *vms1*Δ *rqc2*Δ strain expressing Rqc2p-HA and Vms1p-V5, Vms1p-V5, Rqc2p-HA or the Rqc2p CAT-tailing-defective mutant Rqc2p-D98Y from top to bottom, respectively. Strains were treated with CHX prior to fractionation by sucrose density centrifugation. Chromatographic analysis (*A*_254_) was used to determine the distribution of the 40S, 60S, 80S and polysome content of the 16 collected fractions. Immunoblot analysis was performed only on fractions 3-11. The distribution of the 60S subunit was confirmed by immunoblotting of the ribosomal subunit, Rpl3p. Immunoblotting of HA and V5 was used to detect Rqc2p and Vms1p, respectively. (f) Polysome profiles of whole cell extracts from the *vms1*Δ *rqc2*Δ strain expressing Rqc2p-HA and Vms1p-V5 under the GAL-inducible promoter after galactose induction for 4, 8 and 16 hr from top to bottom, respectively. Chromatographic analysis (*A*_254_) was used to determine the distribution of the 40S, 60S, 80S and polysome content of the 16 collected fractions. The distribution of the 60S subunit was confirmed by immunoblotting of the ribosomal subunit, Rpl3p. Immunoblotting of HA and V5 was used to detect Rqc2p and Vms1p, respectively.

**Extended Data Figure 2.**
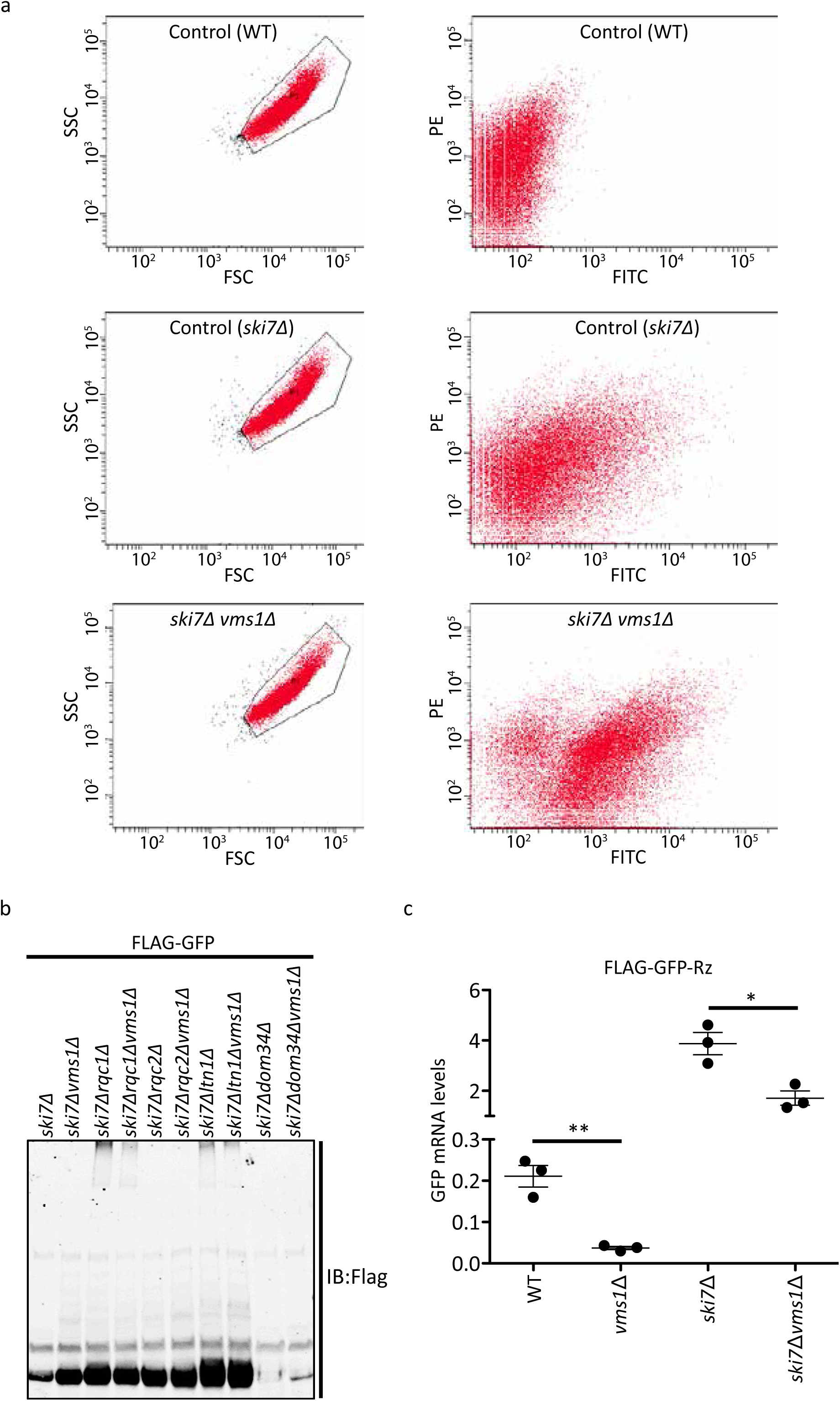
(a) Gating strategy for analyzing FLAG-GFP positive cells. Panel shows gating parameters for collection of total GFP intensity, excluding cellular debris in WT, *ski7*Δ and *ski7*Δ *vms1*Δ. SSC, Side Scatter light; FSC, Forward Scatter light; PE, phycoerythrin; FITC, Fluorescein isothiocyanate. PE was plotted but not analyzed in this study. (b) Immunoblot analysis of whole cell extracts from the indicated strains expressing the FLAG-GFP^Rz^ construct (same as in Fig. 2). Immunoblotting of Flag (overexposed) was used to detect the accumulation of aggregates in the stacking portion of the gel. (c) qRT-PCR analysis of the indicated strains expressing the FLAG-GFP^Rz^ construct (*n*=3, data are mean ± s.e.m. ^⋆⋆^ *P* < 0.002 and ^⋆^ *P* < 0.01, P-value was calculated using unpaired Student’s t-test).

**Extended Data Figure 3.**
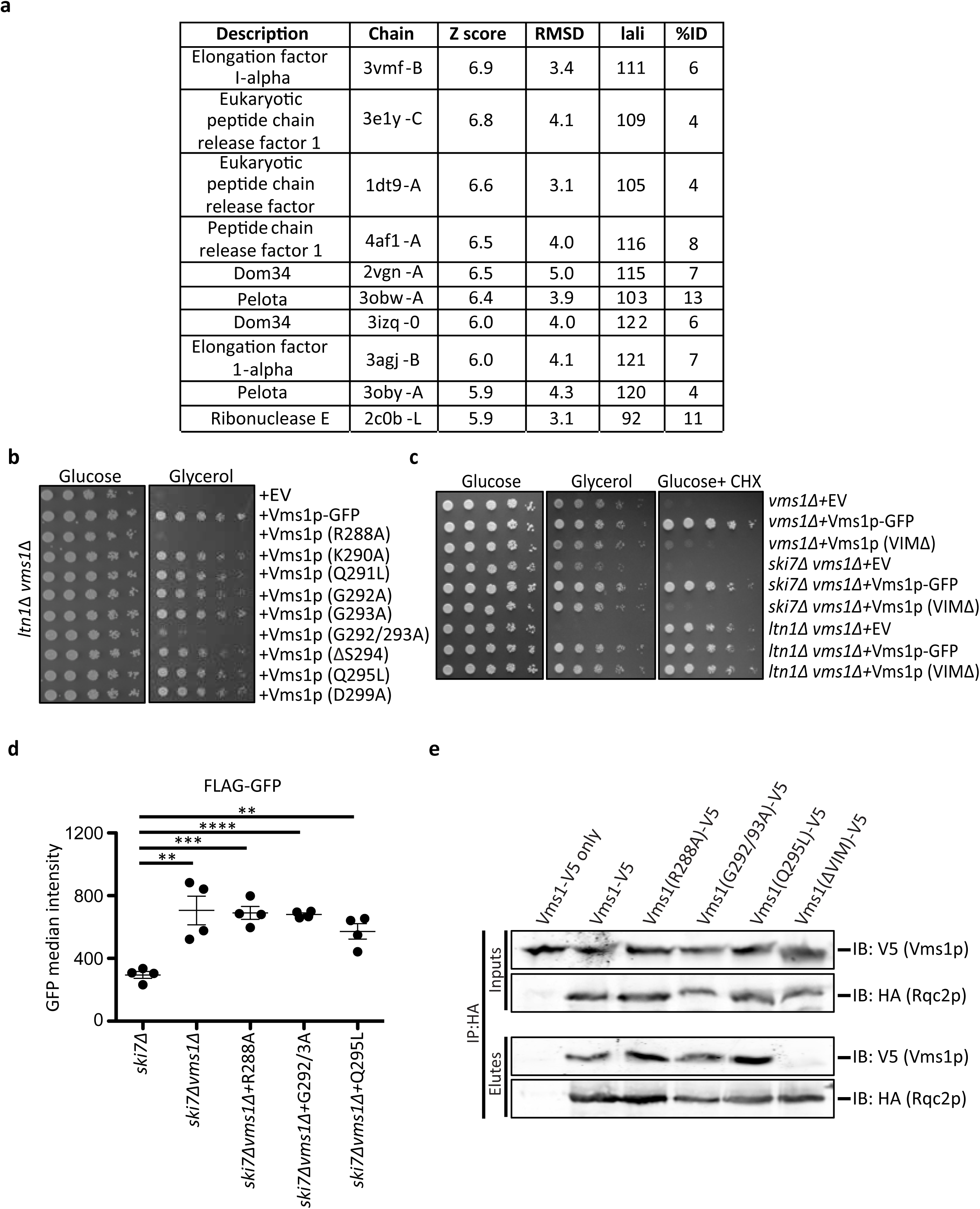
(a) Similar structures to the Vms1p MTD/eRFL returned from the Dali server^27^. Z-score indicates degree of structural similarity, with above 2 being a similar fold. lali, number of aligned residues; %ID, percent identical residues. (b) Serial dilutions of *ltn1*Δ *vms1*Δ cells with the indicated plasmids were spotted on synthetic media supplemented with glucose or glycerol. (c) Serial dilutions of indicated strains were spotted on glucose, glycerol and glucose supplemented with cycloheximide (CHX) and grown for 2 or 3 days, respectively. (d) Flow cytometry quantifications of FLAG-GFP accumulation in the indicated strains. Median GFP intensity values (*n*=4, data are mean ± s.e.m. ^⋆⋆^ *P* < 0.004, ^⋆⋆⋆^ *P* < 0.0002, ^⋆⋆⋆⋆^ *P* < 0.0001, *P*-value was calculated using unpaired Student’s t-test). (e) Immunoprecipitation using the anti-HA antibody in the *rqc2*Δ *vms1*Δ strain expressing Rqc2p and Vms1p-V5 (control); Rqc2p-HA and Vms1p-V5; or Rqc2p-HA and Vms1p-V5 mutants. Immunoblotting of HA and V5 was used to identify Rqc2p and Vms1p, respectively.

**Extended Data Figure 4.**
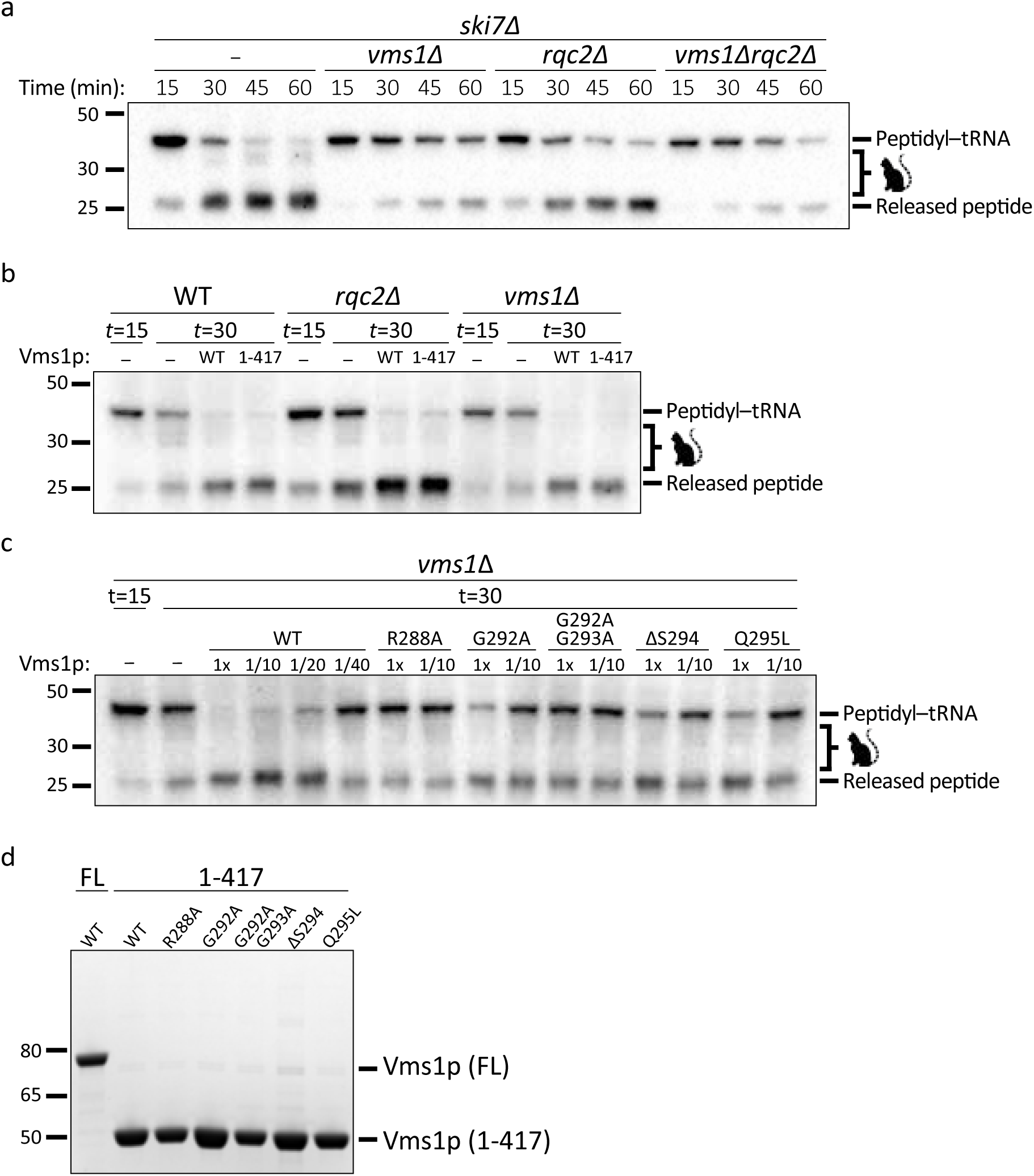
(a) Time courses of *S. cerevisiae* in vitro translation (ScIVT) reactions prepared with a truncated mRNA (lacking a stop codon). Extract genotypes are indicated above. Peptides that have been CAT-tailed and released are denoted by: 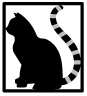 (b) ScIVT reactions prepared as in (a) with WT, *rqc2Δ*, or *vms1Δ* extracts. At *t*=15, buffer (-) or pure protein (4.2 μM final) was added. FL = Full Length Vms1; 1-417 = N-terminus through eRF1-like domain. (c) ScIVT reactions prepared as in (a) with a *vms1Δ* extract. At *t*=15, buffer, WT(1-417), or mutant(1-417) protein was added (see Methods). (d) Coomassie staining of purified Vms1 proteins used in ScIVT rescue experiments. FL = Full Length; 1-417 = N-terminus through eRF1-like domain.

